# Acute systemic macrophage depletion in osteoarthritic mice alleviates pain-related behaviors and does not affect joint damage

**DOI:** 10.1101/2024.08.16.608301

**Authors:** Terese Geraghty, Shingo Ishihara, Alia M. Obeidat, Natalie S. Adamczyk, Rahel S. Hunter, Jun Li, Lai Wang, Hoomin Lee, Frank C. Ko, Anne-Marie Malfait, Rachel E. Miller

## Abstract

**Background:** Osteoarthritis (OA) is a painful degenerative joint disease and a leading source of years lived with disability globally due to inadequate treatment options. Neuroimmune interactions reportedly contribute to OA pain pathogenesis. Notably, in rodents, macrophages in the DRG are associated with onset of persistent OA pain. Our objective was to determine the effects of acute systemic macrophage depletion on pain-related behaviors and joint damage using surgical mouse models in both sexes.

**Methods:** We depleted CSF1R+ macrophages by treating male macrophage Fas-induced apoptosis (MaFIA) transgenic mice 8-or 16-weeks *post* destabilization of the medial meniscus (DMM) with AP20187 or vehicle control (10 mg/kg *i.p.*, 1x/day for 5 days), or treating female MaFIA mice 12 weeks *post* partial meniscectomy (PMX) with AP20187 or vehicle control. We measured pain-related behaviors 1-3 days before and after depletion, and, 3-4 days after the last injection we examined joint histopathology and performed flow cytometry of the dorsal root ganglia (DRGs). In a separate cohort of male 8-week DMM mice or age-matched naïve vehicle controls, we conducted DRG bulk RNA-sequencing analyses after the 5-day vehicle or AP20187 treatment.

**Results:** Eight-and 16-weeks *post* DMM in male mice, AP20187-induced macrophage depletion resulted in attenuated mechanical allodynia and knee hyperalgesia. Female mice showed alleviation of mechanical allodynia, knee hyperalgesia, and weight bearing deficits after macrophage depletion at 12 weeks *post* PMX. Macrophage depletion did not affect the degree of cartilage degeneration, osteophyte width, or synovitis in either sex. Flow cytometry of the DRG revealed that macrophages and neutrophils were reduced after AP20187 treatment. In addition, in the DRG, only MHCII+ M1-like macrophages were significantly decreased, while CD163+MHCII-M2-like macrophages were not affected in both sexes. DRG bulk RNA-seq revealed that *Cxcl10* and *Il1b* were upregulated with DMM surgery compared to naïve mice, and downregulated in DMM after acute macrophage depletion.

**Conclusions:** Acute systemic macrophage depletion reduced the levels of pro-inflammatory macrophages in the DRG and alleviated pain-related behaviors in established surgically induced OA in mice of both sexes, without affecting joint damage. Overall, these studies provide insight into immune cell regulation in the DRG during OA.

## Background

Osteoarthritis (OA) is a major contributor to adult disability and an increasing societal burden, with an estimated 595 million people living with OA worldwide (1). Pain drives OA patients to seek clinical care; yet, current strategies for OA analgesia do not provide adequate pain relief (2).

The role of synovial inflammation in OA pathology has been recognized for some time (3), and a number of studies have shown a positive correlation between synovial inflammation and pain (4, 5). More recently, a link between OA synovitis and sensitization of the nervous system was also identified (6). Of the various types of immune cells that have been found in the OA synovium, macrophages represent the predominant cell type (7), and a novel imaging approach demonstrated that numbers of activated macrophages in OA knees positively correlated with pain severity (8). In addition, elevated levels of soluble macrophage markers in both the synovial fluid and in the blood have been associated with symptom severity as well as disease progression (9, 10), which could be due to the fact that macrophages are able to produce a number of pro-algesic factors, including cytokines and chemokines, that can signal to nociceptors to promote sensitization (11).

As joint-innervating nociceptors become activated, long-term changes can be enacted at the level of the dorsal root ganglia (DRG), where the cell bodies of these neurons are located. We and others have shown that chemokines produced in the DRG can in turn promote the accumulation of macrophages at this site (12–15), in addition to the immune cell changes within the joint itself. Preclinical studies have provided evidence that targeting macrophages in the DRG may be sufficient to provide analgesia. For instance, in a model of nerve injury, depletion of macrophages at the site of nerve injury did not abate pain, while systemic depletion reduced macrophages in the DRG and pain-related behaviors in both male and female mice (16). In addition, in the mouse or rat monoiodoacetate (MIA) model of joint pain, systemic ablation of macrophages reduced pain-related behaviors (13, 17). However, each study only focused on characterizing the effects of systemic depletion in one anatomical location – either in the synovium (17) or the DRG (13), and the effect of depletion on joint damage was not assessed in either of those studies.

While targeting macrophages in the joint and/or in the DRG is potentially beneficial in terms of pain relief, the impact of macrophage depletion on the joint remains unclear. Intra-articular macrophage depletion approaches that began from an early stage were shown to promote synovitis in some models (18–20), while in one, intra-articular depletion of macrophages from an early stage reduced osteophyte formation (21). Here, we aimed to examine the acute effects of systemic macrophage depletion on structural joint changes, pain-related behaviors, and cellular and molecular changes in the DRG in surgically induced OA in mice of both sexes (destabilization of the medial meniscus (DMM) in males, and partial medial meniscectomy (PMX) in females). We focused the intervention on late-stage disease, *i.e.,* time points when both joint structural damage and pain-related behaviors had already developed, in order to provide information on whether targeting macrophages in established OA could be beneficial.

## Methods

### Animals

Macrophage Fas-Induced Apoptosis (MaFIA) mice were purchased from Jackson laboratories (strain name: C57BL/6-Tg(Csf1r-EGFP-NGFR/FKBP1A/TNFRSF6)2Bck/J; RRID:IMSR_JAX:005070) and bred in house (homozygous x homozygous). When male mice became 12 weeks old, destabilization of the medial meniscus (DMM) surgery was performed on right knee joints, as previously described (22). Since young female mice do not develop appreciable joint damage in the DMM model (23), partial meniscectomy surgery (PMX) was performed in the right knee joint of 12-week old female MaFIA mice, as previously described (24). Animals were anesthetized using isoflurane for surgical procedures. Animals were housed at Rush, had unrestricted access to food and water, and were kept on a 12-hour light cycle. All animal experiments were approved by our Institutional Animal Care and Use Committee.

### Macrophage depletion

MaFIA mice were weighed and given 10 mg/kg AP20187 (Tocris, Cat. #6297) or vehicle solution (4% Ethanol, 1.7% Tween-80, 10% PEG-400, water) to achieve macrophage depletion as previously described (25). This otherwise inert pharmacologic agent, AP20187, binds to the CSF1R-eGFP transgenic receptor containing a Fas-cytoplasmic domain, and induces Fas-driven apoptosis in CSF1R+ cells, resulting in systemic macrophage depletion. For male MaFIA mice at 8-weeks *post* DMM surgery, vehicle (n=10) or AP20187 (n=13) was administered intraperitoneally (*i.p.*) once daily for 5 consecutive days, at approximately the same time each day (**Fig. 1**). During the treatment period, mice were housed with softer bedding and dripping water. Vehicle and AP20187 groups were housed in separate cages. Within one to three days before and one to three days after the 5-day depletion period, mice were subjected to pain-related behavior testing. Following the last behavioral test *post* depletion (3-4 days after the last depletion injection), mice were sacrificed for downstream analyses. The same procedure was done in a different cohort of male MaFIA mice at 16 weeks *post* DMM surgery (Vehicle (n=6), AP20187 (n=6)) and in female MaFIA mice at 12 weeks *post* PMX surgery (Vehicle (n=8), AP20187 (n=10)).

**Fig. 1.**
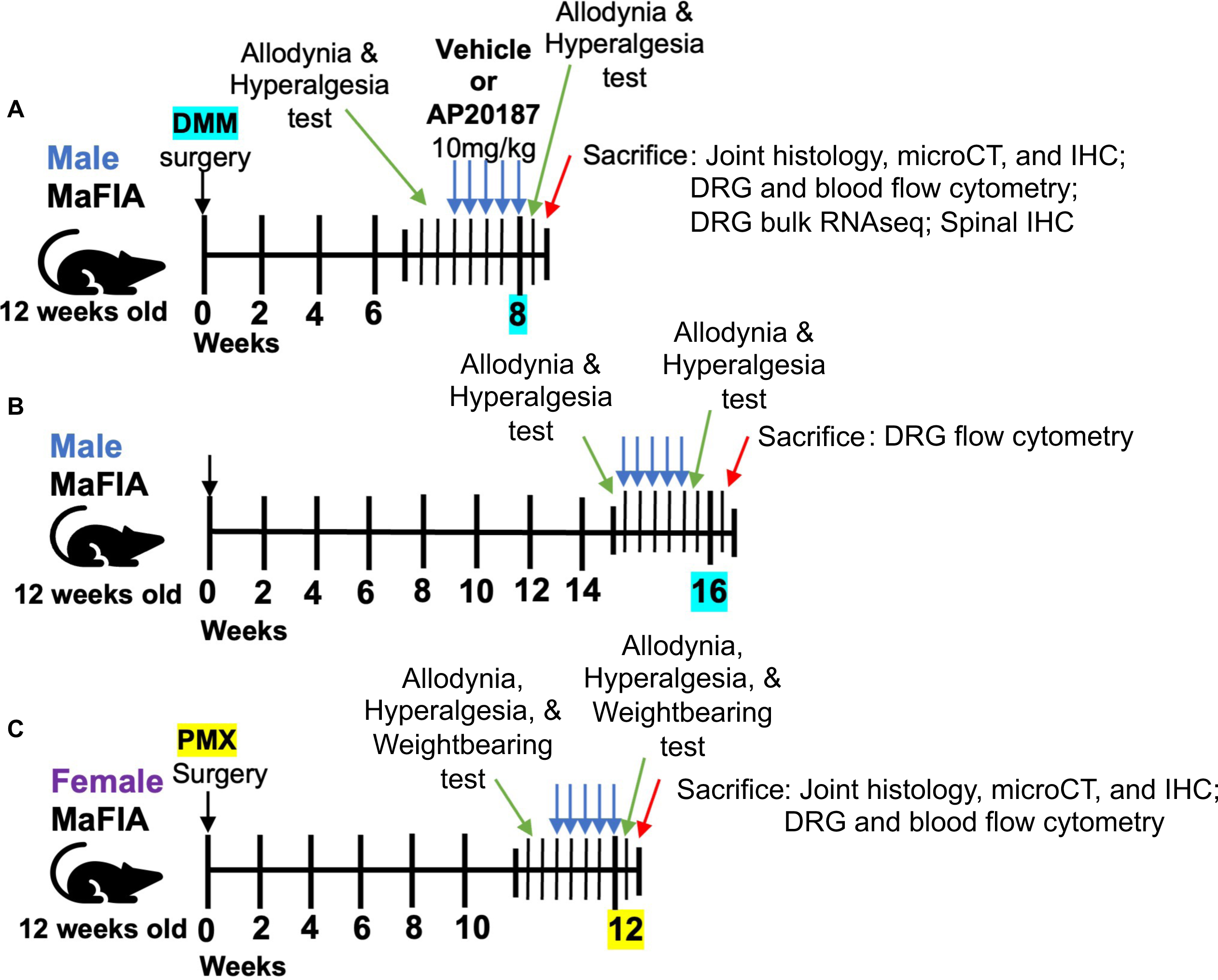
Macrophage depletion experimental design schematic. (A) Male MaFIA mice had DMM surgery at 12 weeks of age. At 7 weeks after DMM, mice were tested for mechanical allodynia and knee hyperalgesia, followed by 5 once daily injections (i.p.) of vehicle solution or AP20187 (10mg/kg). Pain-related behaviors were tested afterward, one test per day. Then mice were sacrificed for downstream analyses. (B) Same as in (A) but vehicle or AP20187 treatment started at 15 weeks after DMM surgery. (C) Female MaFIA mice had PMX surgery at 12 weeks of age and were tested for mechanical allodynia, knee hyperalgesia, and static weightbearing at 11 weeks after PMX. After 5 daily i.p. injections of Vehicle or AP20187, mice were tested for the same pain-related behaviors and then sacrificed for downstream analyses.

### Behavior testing

Male and female MaFIA mice were evaluated for mechanical allodynia and knee hyperalgesia, while weight bearing was tested on female mice only. Only one behavior test was performed per day, and testers were blinded to animal groups. Mice were evaluated before and after the 5-day treatment period.

Mechanical allodynia in the hind paws was measured using von Frey fibers and the 50% withdrawal threshold was calculated via the up-down method, as described previously (12, 26).

Knee hyperalgesia was measured by pressure application measurement (PAM) testing, which applies a range of forces directly to the knee joint to determine a quantitative withdrawal threshold (27, 28). Briefly, mice were restrained by hand and the knee was held in flexion at a similar angle for each mouse. Then, the PAM transducer was pressed against the medial side of the ipsilateral knee and an increasing amount of force was applied up to a maximum of 450g. The force at which the mouse tried to withdraw its knee was recorded. If the mouse did not withdraw, the maximum force of 450g was assigned.

Weight bearing asymmetry was assessed in female MaFIA mice, utilizing a custom voluntarily accessed static incapacitance (VASIC) method where mice were trained to perform a string-pulling task while freely standing on a Bioseb static incapacitance meter platform (Harvard Apparatus) in a custom-built plexiglass chamber (previously described in (29)). To accustom mice to the task, mice were trained one week prior to the first test, as described. Weight bearing was recorded once mice had one hind limb placed on each load cell and were pulling a hanging string to receive a Cheerio reward. Three readings were taken per animal and averaged.

### Joint Histology

Following behavior testing, mice were sacrificed and right-side (ipsilateral-affected) knees were collected for histologic analysis. Knees were formalin fixed, then decalcified in EDTA for 3 weeks, and embedded in paraffin. Six-micron thick sections from the center of the joint were stained with Toluidine Blue or Safranin-O for the evaluation of cartilage damage based on Osteoarthritis Research Society International recommendations (30, 31). These analyses were done semi-quantitatively, and tissue sections were stained altogether when possible. For cartilage degeneration, medial femoral condyles and tibial plateau were scored for severity of cartilage degeneration. For each cartilage surface, scores were assigned individually to each of three zones (inner, middle, outer) on a scale of 0–5, with 5 representing the most damage. The maximum score for the sum of femoral and tibial cartilage degeneration is 30. We also measured osteophyte width, one section with the major osteophyte was assessed using Osteomeasure software (OsteoMetrics) as described (15, 32). The synovial pathology was evaluated as changes in synovial hyperplasia, cellularity, and fibrosis, which were evaluated at the synovial insertion of medial femur and medial tibia separately as described (33). Both joint spaces were visible, except for capsule insertion in some instances, in which the score was considered 0 for that quadrant. Synovitis scoring was performed by 1 independent observer blinded to the groups. Synovial hyperplasia was defined as thickness of the lining layer with a score range of 0-3. Cellularity was defined as the cell density of the synovial sublining with a score range of 0-3. Fibrosis was defined as the extracellular matrix density in the synovium with a score range of 0-1. Each synovial pathology was reported as a sum score of the medial tibial and medial femoral quadrants and reported per knee.

### Joint Microcomputed Tomography

Micro-computed tomographic (µCT) imaging was performed on a subset of intact knee joints using a high-resolution laboratory imaging system (µCT50, Scanco Medical AG, Brüttisellen, Switzerland) in accordance with the American Society of Bone and Mineral Research (ASBMR) guidelines for the use of µCT in rodents (34). Scans were acquired using a 10 µm^3^ isotropic voxel, 70 kVp and 114 µA peak x-ray tube potential and intensity, 500 ms integration time, and were subjected to Gaussian filtration. The subchondral bone analysis was performed by manually contouring the outline of the entire tibial epiphysis. A lower threshold of 375 mg HA/cm^3^ was used to evaluate trabecular bone volume fraction (BV/TV, mm^3^/mm^3^).

### Joint Immunohistochemistry

For TRAP and F4/80 staining, a subset of knee joints were evaluated.

Paraffin sections were used to evaluate osteoclasts in the subchondral bone by enzymatic TRAP staining (35). Within the tibial epiphysis, we counted the total number of osteoclasts (# osteoclasts/mm), which was normalized by bone surface (BS). All measurements were performed using Osteomeasure software (OsteoMetrics, Decatur, GA, USA).

Immunohistochemistry was performed by deparaffinization, antigen retrieval, blocking with normal goat serum, incubating with primary anti-mouse F4/80 antibody (1:100, Abcam ab6640, RRID:AB_1140040) overnight, and developed with DAB staining the next day. Images were taken at 4x and 10x. For quantification, F4/80 images (at 10x) of the medial femoral, medial meniscus, and medial tibial synovium were input into FIJI/Image J (version 1.0). A threshold was set for positive signal detection, and number of regions of interest (ROIs) were counted. For each sample, the ROIs for all three locations in the medial synovium were summed.

### Spinal Immunohistochemistry

For spinal column evaluation the spinal column was collected in a subset of animals. This was done by hydrophobic exclusion, *i.e.,* flushing the intact spine (vertebrae intact), with phosphate buffered saline (PBS), placing the spine in PFA overnight, then 30% sucrose overnight-3 days at 4°C, and embedding in OCT. Upper lumber spinal column was cryosectioned at 10um and stained for chicken anti-mouse Green Fluorescent Protein (GFP) (Abcam) and DAPI (nuclei dye stain, Sigma). CSF1R-GFP+ cells were manually counted in 5 separate areas of the spinal column from 60x images for Vehicle (n=3) or AP20187 (n=3) male MaFIA mice 8 weeks *post* DMM or from Vehicle-treated (n=3) naïve mice as controls.

### Flow Cytometry

Ipsilateral L3-5 DRGs were dissected from DMM or PMX mice and pooled from two mice (3 ipsilateral affected DRGs per mouse = total 6 DRGs), *i.e.,* when the mouse number n=10, flow cytometry sample number n=5. To yield at least 1 million cells, pooling of 6 DRGs was necessary (15). Ipsilateral lumbar levels 3-5 DRG were selected because these are the knee and hind-paw innervating DRGs (36). After dissection, DRGs were digested with collagenase type IV (1.6 mg/mL) and DNase I (200 μg/mL) shaking for 1 hour at 37°C. Following digestion, cells were counted, and 1 million cells were stained with an immune cell panel of anti-mouse antibodies: PE-CD45, AF700-CD3, BV711-CD11b, PE/Cy7-MHCII, PerCP/Cy5.5-Ly6G, APC-F4/80, BV421-CD163, BV605-CCR2 (BioLegend), endogenous signal for GFP-CSF1R, and Aqua-Live/Dead stain (ThermoFisher). After staining, sample data were acquired through an LSR Fortessa flow cytometer. For peripheral blood, additional antibodies were used for T cells, BV421-CD3, AF700-CD8, and PE-Cy7-CD4, and tissue macrophage markers F4/80, MHCII, and CD163 were excluded. Stained sample data were collected through the LSR Fortessa and analyzed by FlowJo software (version 10). See **Supplementary Table 1** for detailed information on flow cytometry antibodies.

### DRG bulk RNA-sequencing

A separate cohort of male MaFIA mice that underwent DMM surgery were treated 8 weeks after surgery with either vehicle (n=5) or AP20187 (n=5), as described above. Age-matched naïve mice (20-weeks old) were treated with vehicle (n=5). After 5 consecutive days of treatment, mice were sacrificed and L3-5 ipsilateral DRGs were collected and lysed in Trizol. RNA was extracted using an RNeasy micro kit (Qiagen) and sent to LC Sciences for sequencing. RNA Integrity Number must have been greater than 6.5 to continue with Poly(A) RNA sequencing. Two samples had RNA integrity numbers less than 6.5 but were determined to be of sufficient quality based on their Electropherogram profile.

Poly(A) RNA sequencing library was prepared following Illumina’s TruSeq-stranded-mRNA sample preparation protocol. Paired-ended sequencing (150 bp) was performed on Illumina’s NovaSeq 6000 sequencing system (LC Sciences). HISAT2 was used to map reads to the Mus musculus reference genome (https://ftp.ensembl.org/pub/release-107/fasta/mus_musculus/dna/) (37). The mapped reads of each sample were assembled using StringTie with default parameters, and StringTie and ballgown were used to estimate the expression levels of all transcripts and perform expression abundance for mRNAs by calculating FPKM (fragment per kilobase of transcript per million mapped reads) value. mRNA differential expression analysis was performed by R package DESeq2 (38) between two different groups (DMM AP *vs*. DMM vehicle; DMM vehicle *vs*. Naïve vehicle). The raw sequence data and counts matrices have been submitted to NCBI Gene Expression Omnibus (GEO) with accession number GSE246252 and the results of the differential expression analysis are provided in **Supplemental Tables 2and 3**. The focus of our analysis was two-fold: 1) to identify genes related to immune cell function that were expressed at levels above FPKM>1 and regulated in the DRG in the current study (DMM-vehicle *vs*. naïve-vehicle) as well as differentially regulated in other published DRG datasets from OA models (14, 39, 40); and 2) to identify whether any of these genes were regulated after macrophage depletion (DMM-AP *vs*. DMM-vehicle).

### Statistical analysis

Statistical calculations were performed using GraphPad Prism 9. Paired two-tailed t-tests were used for before and after macrophage depletion comparison for MaFIA mice pain-related behavior studies. For mechanical allodynia, since the von Frey fibers are on a log scale, the data was first log-transformed and then a paired student’s t test was used for pairwise comparison. Unpaired two-tailed student’s t-test was used for histology analysis, except for synovitis scores, where Mann-Whitney test was used. For flow cytometry, Mann-Whitney test was used. Unpaired two-tailed t tests were used in all other analyses. Mean ± 95% Confidence Interval (CI) is shown in all graphs and p values are stated on each graph. P values were considered significant if less than 0.05.

## Results

### Systemic macrophage depletion reduces pain-related behaviors in both sexes

To determine if the acute depletion of macrophages in the DRG changes pain-related behaviors in established OA, we utilized MaFIA transgenic mice (25) to perform conditional macrophage depletion at time points after surgery when mice had developed joint damage and pain-related behaviors. In brief, we tested mechanical allodynia and knee hyperalgesia in male MaFIA mice 7 weeks after DMM, followed by 5 consecutive days of treatment with AP20187 (10 mg/kg *i.p.*) or vehicle (**Fig. 1A**). We confirmed macrophage depletion when mice were taken down following the last behavioral test, 3 days after the last injection, by evaluating CSF1R-GFP+ cells by flow cytometry of DRGs (**Suppl. Fig. 1**) and peripheral blood (**Suppl. Fig. 2**). We also checked if AP20187 depletion affected CSF1R+ cells in the dorsal horn of the spinal cord, and found no difference in GFP+CSF1R+ cells in the lumbar dorsal horn between AP20187 and vehicle-treated male DMM mice (**Suppl. Fig. 3**). This suggests acute macrophage depletion has no effect on CSF1R+ microglia in the central nervous system, as others have reported (16).

Immediately after macrophage depletion, osteoarthritic mice, 8 weeks *post* DMM, had significantly improved mechanical allodynia (**Fig. 2A**) and knee hyperalgesia (**Fig. 2B**). To determine if macrophage depletion is also analgesic in late-stage OA, we used a new cohort of male mice and depleted macrophages 15 weeks *post* DMM (**Fig. 1B**). Again, we found significant alleviation of mechanical allodynia (**Fig. 2C**) and knee hyperalgesia (**Fig. 2D**).

**Fig. 2.**
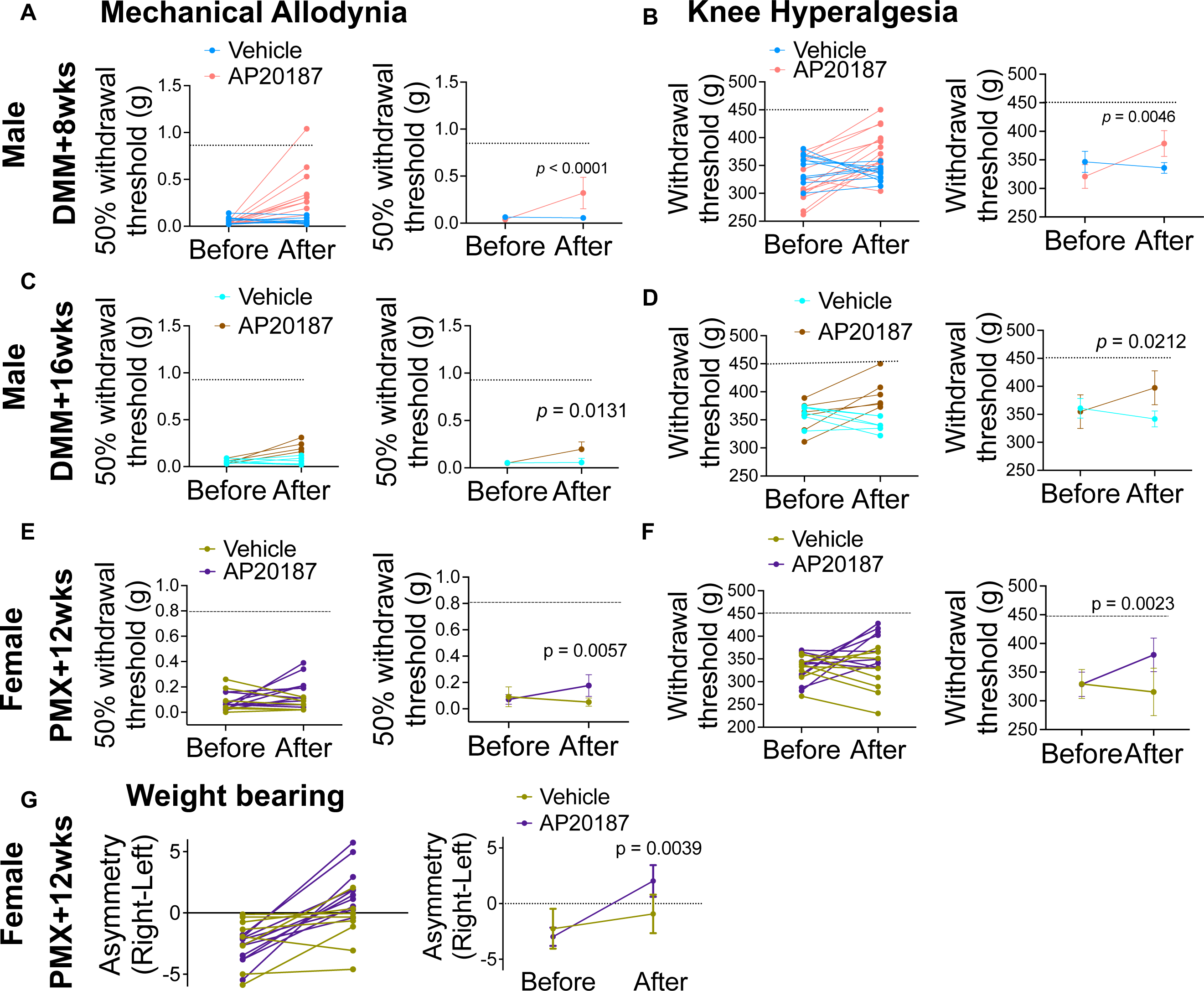
Macrophage depletion alleviated pain-related behaviors in mice of both sexes with OA. (A) Mechanical allodynia and (B) Knee hyperalgesia withdrawal thresholds in male MaFIA mice at 8 weeks after DMM surgery showing before and after treatment with Vehicle (in blue, n=10) or AP20187 (in red, n=13). The left graph shows individual mouse data points before and after treatment, and the right graph shows the mean +/-95% CI of each group’s withdrawal thresholds before and after treatment. (C) and (D) Same as in (A) and (B) but tested male mice at 16 weeks after DMM surgery; before and after Vehicle (in aqua, n=3) or AP20187 (in brown, n=3) treatment. (E) Mechanical allodynia and (F) Knee hyperalgesia withdrawal thresholds in female MaFIA mice at 12 weeks after PMX surgery showing before and after Vehicle (in dark yellow, n=8) or AP20187 (in purple, n=10) treatment. (G) Static weight bearing in same cohort of female MaFIA mice at 12 weeks after PMX surgery showing before and after Vehicle (in dark yellow, n=8) or AP20187 (in purple, n=10) treatment. Statistical analysis by paired t-test. P values stated on graphs; considered significant if p < 0.05.

We next asked if macrophage depletion would have the same effect in female mice with experimental OA. We performed PMX surgery in 12-week old female MaFIA mice, and depleted macrophages 11 weeks *post* PMX (**Fig. 1C**). This resulted in significantly attenuated mechanical allodynia (**Fig. 2E**), knee hyperalgesia (**Fig. 2F**). In female mice, we also assessed weightbearing deficits and found that macrophage depletion resulted in improved weight bearing on the ipsilateral knee (**Fig. 2G**) in AP20187-treated *vs*. vehicle-treated mice.

### Acute macrophage depletion effects in the joint

We checked F4/80+ macrophage levels in the synovium in mice 8 weeks *post* DMM (3 days after the end of depletion) and found a significant decrease in F4/80+ synovial macrophages in knees from AP20187 *vs.* vehicle-treated mice (**Suppl. Fig. 4A, B**). Since CSF1R is reportedly expressed on osteoclasts (41), we evaluated if osteoclasts were affected by AP20187 treatment by staining for tartrate-resistant acid phosphatase (TRAP) in mice of both sexes. In addition, we evaluated whether depletion affected overall bone mass using micro-CT, because one study also reported changes in bone morphology within one week of depletion (18). Eight weeks *post* DMM in male mice, acute macrophage depletion did not change the number of TRAP+ osteoclasts (**Suppl. Fig. 4C,D**) nor the bone volume to trabecular volume (BV/TV) ratio (**Suppl. Fig. 4E**). Likewise, we found no difference in the number of TRAP+ osteoclasts (**Suppl. Fig. 4F,G**) or in BV/TV (**Suppl. Fig. 4H**) 12 weeks *post* PMX in female mice.

We next examined cartilage degeneration and synovitis scores. As seen in representative images of knee histopathology at 8-weeks *post* DMM from vehicle or AP20187 treated male mice (**Fig. 3A**), macrophage depletion did not affect the severity of medial cartilage degeneration (**Fig. 3B**), osteophyte width (**Fig. 3C**), or synovitis (**Fig. 3D**), including no difference in medial synovial hyperplasia (**Fig. 3E**), cellularity (**Fig. 3F**), or fibrosis (**Fig. 3G**). We found the same lack effect in AP20187-treated female mice 12 weeks after PMX surgery compared to vehicle (**Fig. 3H**): there was no difference in medial cartilage degeneration (**Fig. 3I**), osteophyte width (**Fig. 3J**), or synovitis (**Fig. 3K**), including no difference in medial synovial hyperplasia (**Fig. 3L**), cellularity (**Fig. 3M**), or fibrosis (**Fig. 3N**). These data suggest acute macrophage depletion did not affect subchondral bone mass, the degree of cartilage degeneration, or synovitis in male or female mice with established OA.

**Fig. 3.**
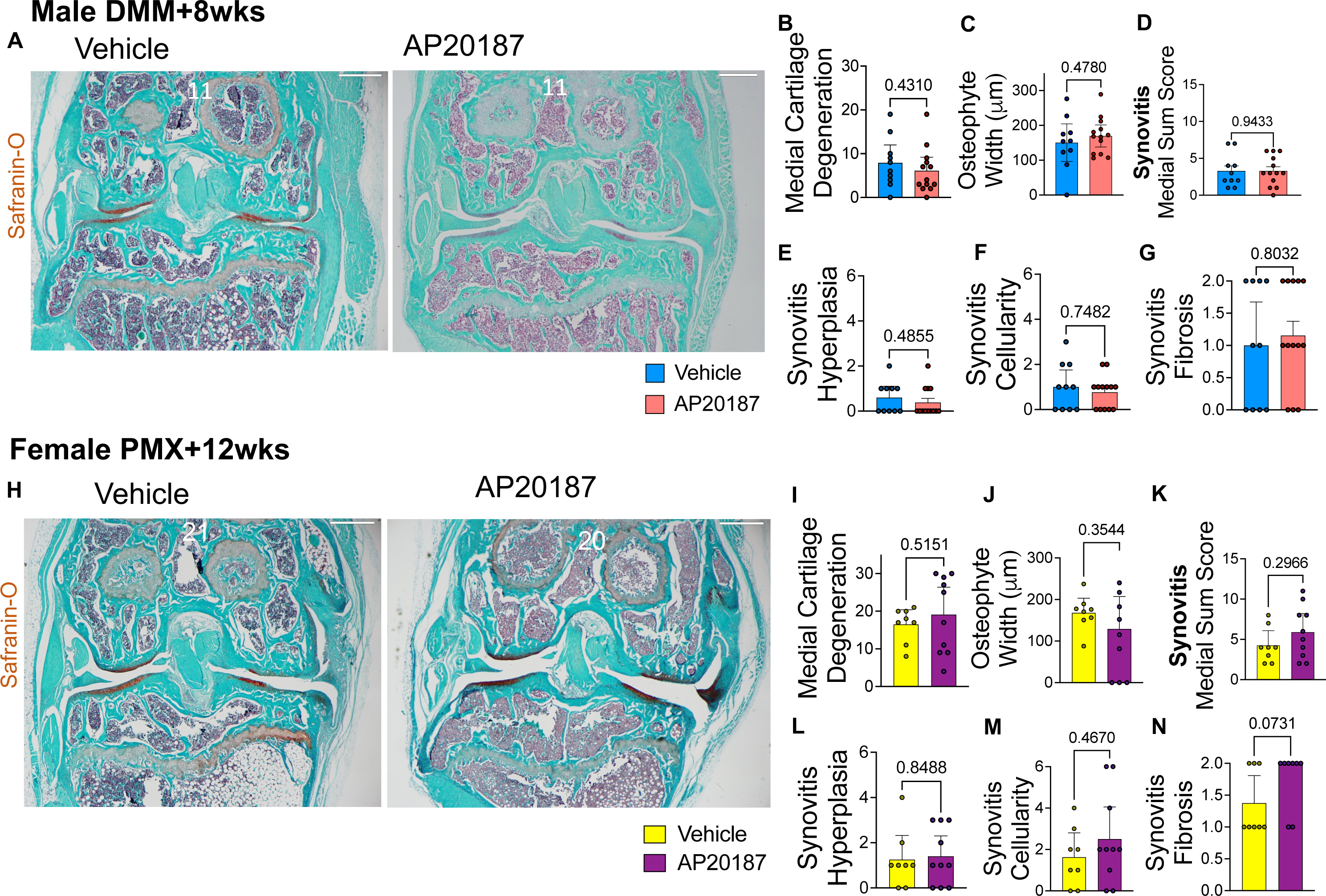
Acute systemic macrophage depletion does not affect joint damage severity. (A) Representative Safranin-O stained Vehicle or AP20187-treated male MaFIA mouse knee images at 8 weeks after DMM surgery; 2x magnification. White numbers represent the cartilage degeneration score for that mouse. (B) Medial cartilage degeneration, (C) Osteophyte width (micrometers), (D) Synovitis medial sum score, (E) Synovial medial hyperplasia, (F) Synovial medial cellularity, and (G) Synovial medial fibrosis scored for male MaFIA mice at 8 weeks after DMM surgery from Vehicle (in blue, n=10) and AP20187 (in red, n=13) groups. (H) Representative Safranin-O stained Vehicle or AP20187-treated female MaFIA mouse knee images at 12 weeks after PMX surgery; 2x magnification. White numbers represent cartilage degeneration score for that mouse. (I) – (N) Same as in (B) – (G) but showing scores for female MaFIA mice at 12 weeks after surgery from Vehicle (in yellow, n=8) or AP20187 (in purple, n=10) treated mice. Scale bar is 500 μm. Statistical analysis for cartilage degeneration and osteophyte width by two-tailed unpaired t-test, and for synovitis scoring, Mann-Whitney test. Graphs show mean +/-95% CI. P values stated on graphs; considered significant if p < 0.05.

### CSF1R+ cells and neutrophils are reduced in the DRG in both sexes after macrophage depletion

To determine the immune cell changes in the DRG with and without macrophage depletion we used flow cytometry to examine DRG immune cell phenotypes. The representative DRG gating strategy is shown in **Suppl. Fig. 1A**. In DRGs from male mice, there was no change in the frequency of CD45+ leukocytes (**Fig. 4A**), CD11b+ myeloid cells (**Fig. 4B**), or CD3+ T cells (**Fig. 4C**) between vehicle and AP20187 treated mice at 8- or 16-weeks *post* DMM. As expected, CSF1R+ (**Fig. 4D, Suppl. Fig. 1B)** monocyte/macrophages were decreased at both timepoints *post* DMM. Ly6G+ neutrophils (**Fig. 4E**) were also significantly decreased after macrophage depletion at 8- weeks, but not 16-weeks *post* DMM surgery, though there was a trending decrease at this time point.

**Fig. 4.**
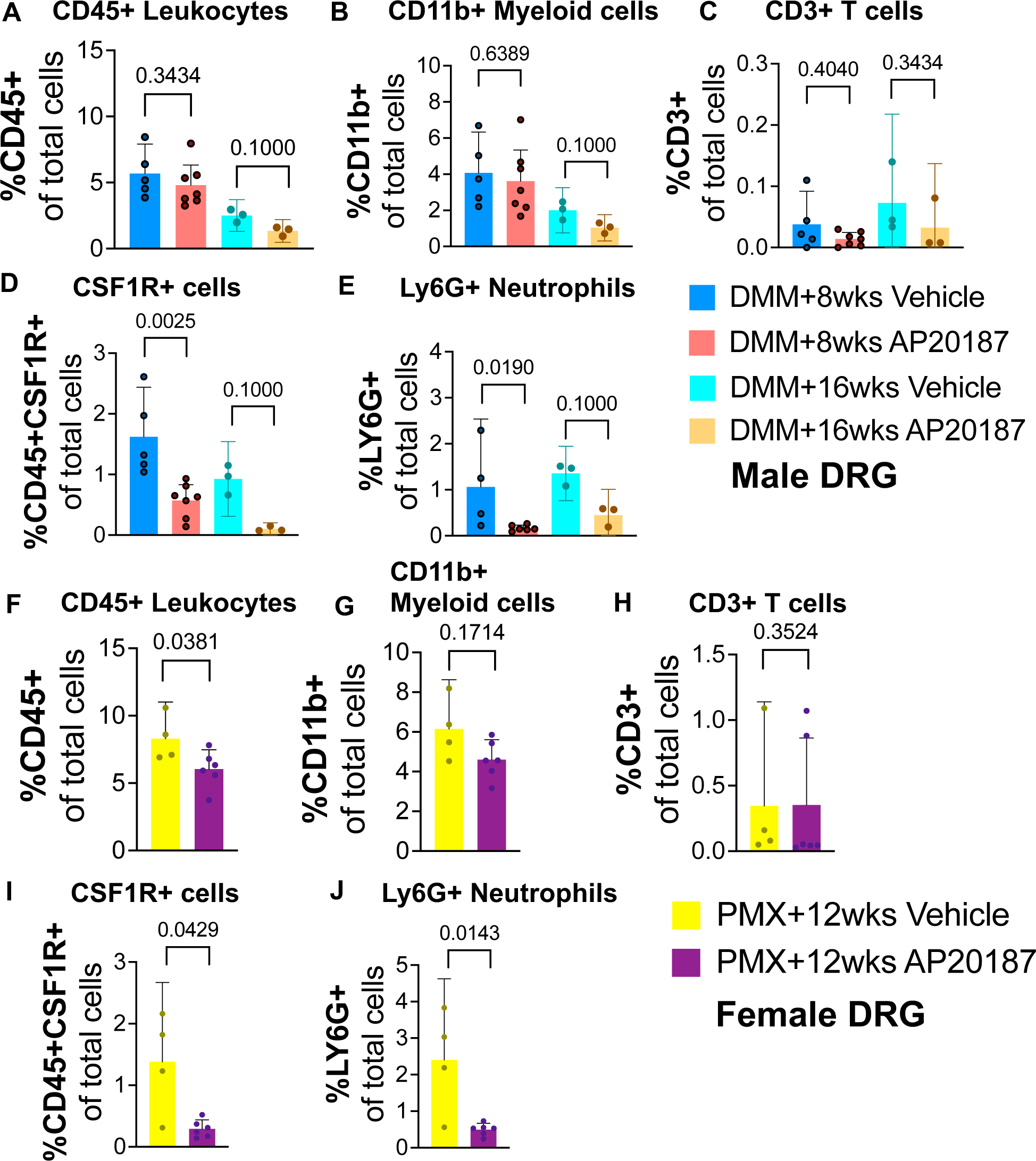
Decreased Ly6G+ neutrophil and CSF1R+ macrophage populations in the DRG after macrophage depletion. Frequency of (A) CD45+ leukocytes, (B) CD11b+ myeloid cells, (C) CD3+ T cells, (D) CSF1R+ monocyte/macrophages, (E) Ly6G+ neutrophils in the DRG of male MaFIA mice at 8 weeks after DMM surgery for Vehicle (blue, n=5) or AP20187 (red, n=7), or at 16 weeks after DMM surgery for Vehicle (teal, n=3) or AP20187 (brown, n=3) treated mice. (F) – (J) Same as in (A) – (E) but showing frequencies of each cell population for female MaFIA mice at 12 weeks after surgery for Vehicle (yellow, n=4) or AP20187 (purple, n=5) treated mice. Statistical analysis by Mann-Whitney test. Error bars show mean +/-95% CI. P values stated on graphs; significant if p < 0.05.

For female DRGs, there was a significant decrease in CD45+ leukocytes (**Fig. 4F)**, and no difference in CD11b+ myeloid cells (**Fig. 4G**) or CD3+ T cells (**Suppl. Fig. 4H**) after macrophage depletion with AP20187 compared to vehicle, treated 12 weeks *post* PMX. CSF1R+ monocyte/macrophages (**Fig. 4I, Suppl. Fig. 1C**) and Ly6G+ neutrophils (**Fig. 4J**) were significantly decreased after macrophage depletion.

Altogether, this shows that AP20187 treatment reduced neutrophils and CSF1R+ cells (macrophages) as well as neutrophils in the DRG in both males and female with experimental OA, while overall CD45+ leukocytes were decreased in females only.

### Effect of macrophage depletion on DRG macrophage phenotypes

Finally, we aimed to characterize the functional phenotypes of DRG macrophages by evaluating the M1-like activated/pro-inflammatory MHCII+ macrophages and M2-like anti-inflammatory CD163+ macrophages. There was a trending decrease in F4/80+ macrophages in AP20187-treated mice compared to vehicle in males at both timepoints *post* DMM (**Fig. 5A**). Under F4/80+ macrophages, we first looked at total F4/80+MHCII+ macrophages and F4/80+CD163+ macrophages. We then made quadrants of MHCII and CD163 positive and negative populations, resulting in CD163+MHCII-, CD163+MHCII+, CD163-MHCII-, and CD163-MHCII+ macrophage populations. In male DRGs, total MHCII+ macrophages were significantly decreased at 8-weeks *post* DMM (**Fig. 5B**), while total F4/80+CD163+ macrophages did not change at either the 8- or 16-week timepoint (**Fig. 5C**). There were also no differences in CD163-MHCII-(**Fig. 5D**) or CD163+MHCII-(**Fig. 5G**) at either timepoint. However, there was a significant decrease in the CD163-MHCII+ (**Fig. 5E**) and CD163+MHCII+ (**Fig. 5F**) populations at 8-weeks *post* DMM in AP20187-treated *vs*. vehicle mice. This suggests a stronger decrease in MHCII+ M1-like macrophages than CD163+ M2-like macrophages in DMM male mice DRGs. In female DRGs, similar to males, there was a trending decrease in total F4/80+ macrophages in the DRG in AP20187-treated mice compared to vehicle (**Fig. 5H**). Total MHCII+ macrophages were significantly decreased at 12-weeks *post* PMX (**Fig. 5I**) and total F4/80+CD163+ macrophages did not change (**Fig. 5J**). There was no difference in CD163-MHCII-(**Fig. 5K**) or CD163+MHCII-(**Fig. 5N**) populations, and a significant decrease in CD163-MHCII+ (**Fig. 5L**) and CD163+MHCII+ (**Fig. 5M**) populations following depletion. This suggests, just like in males, that macrophage depletion with AP20187 compared to vehicle results in a significant decrease in MHCII+ M1-like macrophages with little to no effect on CD163+ M2-like macrophages.

**Fig. 5.**
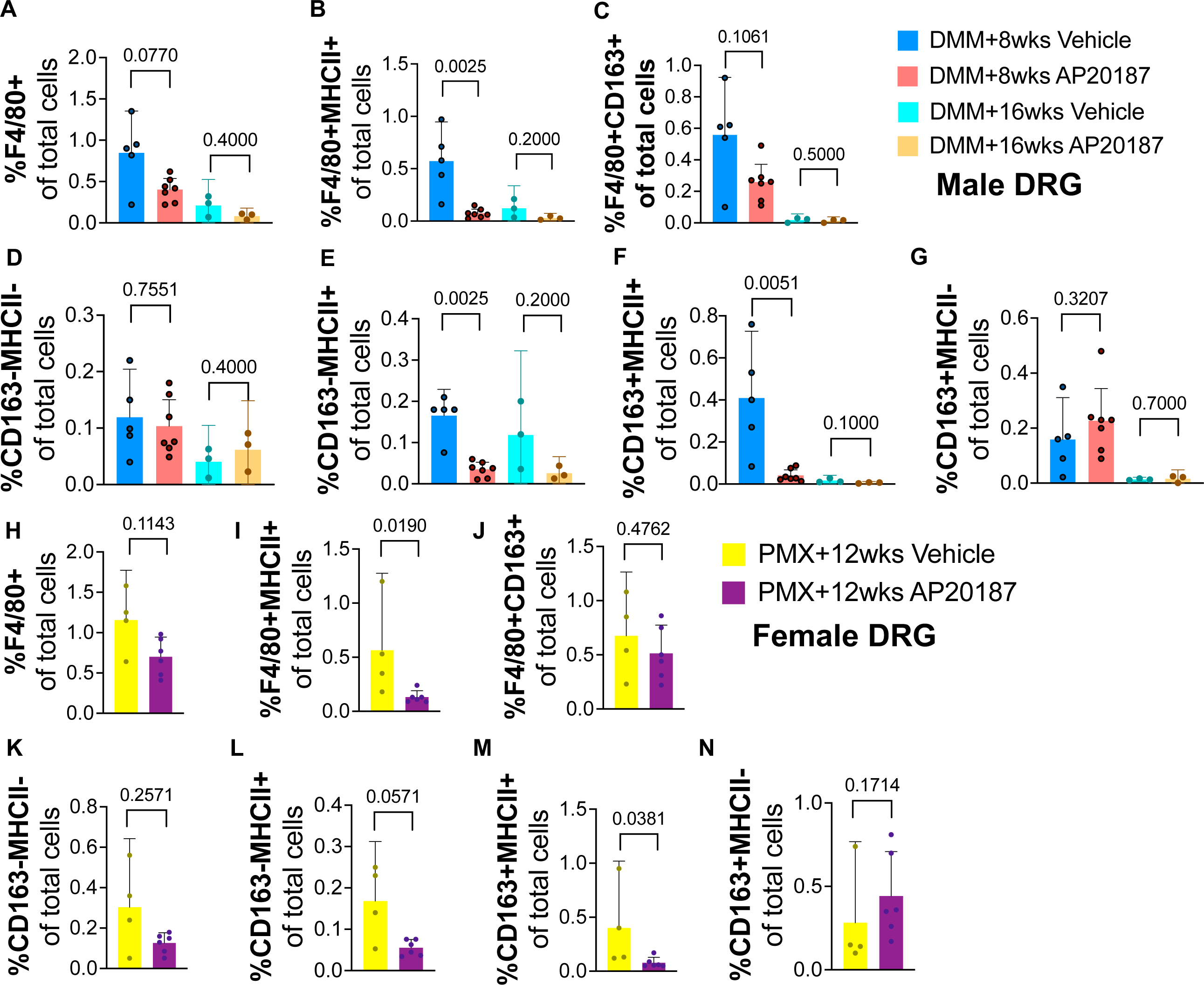
Decreased MHCII+ macrophages, but no change in CD163+ macrophages in the DRG after systemic macrophage depletion in both sexes. (A) Frequency of total F4/80+ macrophages, (B) F4/80+MHCII+, (C) F4/80+CD163+, (D) CD163-MHCII-, (E) CD163-MHCII+, (F) CD163+MHCII+, (G) CD163+MHCII-macrophage populations in the DRG in male MaFIA mice at 8 weeks after DMM surgery for Vehicle (blue, n=5) or AP20187 (red, n=7), or at 16 weeks after DMM surgery for Vehicle (teal, n=3) or AP20187 (brown, n=3) treated mice. (H) – (N) Same as in (A) – (G) but showing frequencies of each cell population for female MaFIA mice at 12 weeks after surgery for Vehicle (yellow, n=4) or AP20187 (purple, n=5) treated mice. Statistical analysis by Mann-Whitney test. Error bars show mean +/-95% CI. P values stated on graphs; significant if p < 0.05.

### Effect of macrophage depletion on DRG gene expression

To understand the molecular changes in the macrophage-depleted DRG, we conducted bulk RNA-sequencing analyses of DRGs from male mice 8-weeks after DMM surgery, with vehicle or with AP20187 treatment, and age-matched naïve controls treated with vehicle. We selected a group of immune-related genes to focus on based on DRG molecular studies from the literature (12, 15, 40, 42–46). Here, in DMM mice treated with vehicle compared to vehicle-treated naïve mice, we found *Ccl2, Ccl3, Ccl7, Cd200r1, Cx3cr1, Cxcl10, Il1b, Il25, Il36g, S100a8, S100a9,* and *Tlr2,* were all upregulated compared to naïve (**Table 1**, DMM-veh *vs*. naïve-veh). We also compared our results to other available published datasets (14, 39, 40), and looked for genes consistently regulated in at least 2 of the 5 OA datasets (4 published datasets or the newly presented DMM-veh *vs*. naïve-veh dataset here) (**Table 1**). Across the different datasets, *Ccl2*, *Ccl3*, *Ccr5*, *Cx3cr1*, *Cd200r1*, *Il1b* were consistently upregulated in the DRG with OA.

**Table 1.** Regulation of genes related to immune cell function in the dorsal root ganglia (DRG) in mouse models of OA.

| Gene | From Ref (14):<br>Early DMM<br>microarray<br>(fold change) | From Ref (14):<br>Persistent<br>DMM<br>microarray<br>(fold change) | From Ref (40): MIA<br>model<br>qPCR (fold<br>change) | From Ref (39): MJL vs<br>unloaded<br>microarray<br>(fold change) | This Study:<br>DMM veh vs<br>Naïve veh<br>Bulk<br>RNAseq +8<br>weeks<br>(log2(fc)) | This Study:<br>DMM AP vs<br>DMM veh<br>Bulk RNAseq<br>+8 weeks<br>(log2(fc)) |
| --- | --- | --- | --- | --- | --- | --- |
| **Ccl2 (up in OA)** | no change | 1.06 (p=0.06) | 1.4221 | 1.39 (p=0.02) | 1.22 (p=0.01) | 1.14 (p=0.03) |
| **Ccl3 (up in OA)** | no change | no change | 1.5999 | no change | 2.79 (p=0.009) | no change |
| Ccl7 | no change | no change | no change | no change | 1.50 (p=0.01) | 1.14 (p=0.04) |
| Ccl12 | no change | 1.24 (p=0.03) | -1.5605 | no change | no change | 1.54 (p=0.002) |
| **Ccr5 (up in OA)** | 1.17 (p=0.08) | 1.08 (p=0.03) | not measured | no change | no change | 1.83 (p=0.00006) |
| Cx3cl1 | no change | 1.14 (p=0.01) | no change | no change | no change | no change |
| **Cx3cr1 (up in OA)** | no change | 1.26 (p=0.004) | not measured | no change | 1.14 (p=0.03) | no change |
| Cxcl10 | no change | no change | -2.2689 | no change | 0.92 (p=0.07) | -0.97 (p=0.009) |
| Cxcl11 | no change | no change | 7.558 | no change | no change | -1 (p=0.01) |
| Cxcr3 | no change | no change | not measured | no change | no change | no change |
| Cd14 | 1.13 (p=0.04) | no change | not measured | no change | no change | 1.84 (p=0.00005) |
| **Cd200r1 (up in OA)** | no change | 1.03 (p=0.04) | not measured | no change | 1.27 (p=0.02) | 2.28 (p=3e-7) |
| Csf1r | 1.27 (p=0.002) | no change | not measured | no change | no change | no change |
| Il10ra | no change | 1.2 (p=0.002) | not measured | no change | no change | no change |
| Il13ra | no change | 1.13 (p=0.03) | not measured | no change | no change | Il13ra1: 1.16 (p=3e-6) |
| **Il1β (up in OA)** | no change | 1.07 (p=0.09) | 1.5562 | no change | 1.24 (p=0.08) | -1.67 (p=0.03) |
| Il25 | no change | no change | not measured | no change | 1.92 (p=0.0007) | no change |
| Il36g | not present | not present | not measured | not present | 1.38 (p=0.02) | 2.33 (p=0.00002) |
| Spp1 | no change | no change | 1.9697 | -1.2 (p=0.09) | no change | no change |
| S100a8 | no change | no change | not measured | no change | 1.37 (p=0.047) | 2.93 (p=2.5e-6) |
| S100a9 | no change | no change | not measured | no change | 1.31 (p=0.05) | 2.86 (p=2.3e-6) |
| Tlr2 | no change | no change | not measured | no change | 1.00 (p=0.09) | 1.88 (p=0.0001) |

DMM=destabilization of the medial meniscus; MIA = monoiodoacetate; MJL = mechanical joint loading; veh = vehicle; AP = AP20187. Bold genes = genes that changed in the same direction in at least 2 mouse models of osteoarthritis. Blue labeled cells = genes where DMM AP vs DMM veh regulated genes in the opposite direction as DMM veh vs Naïve veh.

After macrophage depletion, chemoattractant genes such as *Ccl2* and *Ccl7* were further upregulated (**Table 1**, DMM-AP *vs*. DMM-veh), presumably as a response to return macrophages to the DRG, while the pro-inflammatory genes *Il1b* and *Cxcl10* were downregulated (**Table 1**, DMM-AP *vs*. DMM-veh), suggesting that the decrease in these genes contributed to the associated reduction in pain-related behaviors.

Finally, we also examined gene expression patterns to determine if M1-like or M2-like macrophage patterns changed with depletion. We found that signature genes expressed by CD163+ macrophages (47) were upregulated (*Cd163*, *Fcrls*, *Mrc1*), while genes expressed by MHCII+ macrophages were downregulated (*Ccr2*, *H2-Aa*) (**Table 2**), supporting the flow cytometry results that suggested that MHCII+ macrophages were reduced with depletion and contributed to pain-related behaviors (**Fig. 5**).

**Table 2.** Macrophage marker genes regulated in the DRG after AP20187 treatment.

|  | Gene | DMM AP vs<br>DMM veh DRG<br>Bulk RNAseq<br>+8 weeks<br>(log2(fc)) |
| --- | --- | --- |
| M2-like | Fcrls | 1.35<br>(p=0.03) |
|  | Cd163 | 1.11<br>(p=0.01) |
|  | Mrc1 | 1.44 (p=0.001) |
| M1-like | Ccr2 | -2.64<br>(p=0.000004) |
|  | H2-Aa | -1.87 (p=0.001) |

### Acute macrophage depletion effects on body weight

As seen previously with systemic macrophage depletion (16, 25, 48), we did observe a significant drop in bodyweight in all groups of OA MaFIA mice treated with AP20187 compared to the respective Vehicle mice, while no difference in bodyweight was observed when AP20187 was administered to WT naïve mice (**Suppl. Fig. 5**).

## Discussion

In summary, targeting macrophages systemically in established experimental OA, when joint damage and pain-related behaviors have developed, resulted in reduced pain-related behaviors in both sexes. We observed no acute adverse effects on joint degeneration, synovitis, or osteophytes. Macrophage depletion reduced MHCII+ M1-like pro-inflammatory macrophages in the DRG, while CD163+MHCII-M2-like macrophage numbers were unchanged. Microglia in the dorsal horn were not acutely affected by this depletion protocol. After examining differentially expressed genes in the DRG in various OA datasets across the literature (14, 39, 40), we confirmed the upregulated expression of several inflammatory chemokines and cytokines in the DRG, but only *Il1b* and *Cxcl10* were reversed as a result of macrophage depletion in male DRGs at 8-weeks *post* DMM. Upregulation of these cytokines in the DRG has been implicated by previous work investigating other types of painful conditions (16, 46, 49–52), and together with the work here suggests these cytokines may play a role in supporting OA persistent pain via DRG macrophages.

Our observation that systemic depletion of CSF1R+ cells attenuated pain-related behaviors is consistent with previous studies in other rodent models of knee OA. In the rat MIA model, an intravenous injection of Clophosome restored grip strength (17), and in the mouse MIA model, systemic ablation of macrophages through diphtheria toxin targeting of CSF1R+ cells reduced hind paw mechanical allodynia and weight bearing asymmetry (13). Repeated intra-articular injection of clodronate liposomes from week 2 after surgery also prevented the development of knee hyperalgesia and hind paw mechanical allodynia in the rat anterior cruciate ligament transection and destabilization of the medial meniscus (ACLT/DMM) surgery model (20). This observation contrasts with a study using a mouse nerve injury model, where only systemic depletion, and not local nerve injury site depletion, of macrophages reduced hind paw mechanical allodynia (16). Together, these results suggest that both targeting DRG and knee macrophages could be beneficial for reducing OA pain-related behaviors.

Because of previous concerns regarding the effects of macrophage depletion on joint pathology (18–20), we examined several different aspects of joint damage in this study. In both male and female mice, systemic depletion of macrophages did not result in worsening of cartilage degeneration, synovitis, osteophyte width, or subchondral bone mass when assessed approximately one week later. However, as others have noted (16, 25, 48), mice did lose a significant percentage of body weight as a result of the depletion. Our findings may have differed from previous work due to a difference in the location and/or timing of depletion. For instance, in a closed fracture model, intra-articular depletion of macrophages 2 days prior to injury resulted in increased synovitis and reduced bone mineral density one week later, whereas depletion at the time of injury had no effect on either measure (18). Similarly, in the rat ACLT/DMM surgery model, repeated intra-articular depletion of synovial macrophages beginning 2 weeks *post* surgery resulted in increased synovial fibrosis, vascularization, and perivascular edema by 12 weeks *post* surgery (20). Finally, when macrophage depletion was performed systemically at the time of surgery in a mouse model of OA induced by high fat diet and DMM surgery, this promoted increased synovitis by 9 weeks *post* depletion (19). In contrast, in the collagenase-induced OA model, intra-articular depletion of macrophages prior to the injection of collagenase reduced synovial inflammation and fibrosis as well as osteophyte formation by days 7 and 14 (21). Similarly, systemic depletion at the time of injury in an annular puncture induced disc herniation mouse model reduced ectopic bone formation in herniated discs and adjacent cortical bones 20 days later (48). The local environment of immune cells changes rapidly following joint injury (53), and thus avoiding critical periods when macrophages are needed for resolution of joint inflammation may at least partially explain the differing effects observed. Here, systemic macrophage depletion was performed at a time point in the model when knee inflammation had already peaked (23, 24, 54, 55), which may explain why we did not observe any worsening of damage with depletion.

We have previously demonstrated that increased numbers of F4/80+ macrophages can be found in the DRG 8-16 weeks *post* DMM (12, 14). Here, the reduction in pain-related behaviors observed following systemic ablation of CSF1R+ cells was associated with a decrease specifically in the Ly6G+ population and in the pro-inflammatory MHCII+ macrophage population in the DRG in both sexes, as well as with downregulated expression of *Il1b*, *Cxcl10*, and M1 marker genes (*Ccr2*, *H2-Aa*) in the DRG. This is consistent with another study, where injection of M1-like macrophages adjacent to the DRG induced mechanical allodynia of the hind paw in naïve mice, while injection of M2-like macrophages reduced mechanical hypersensitivity in the MIA model (13). However, it contrasts with the response in the DRG after systemic ablation in a model of disc herniation where increased numbers of F4/80+ macrophages and increased expression of inflammatory cytokines was observed in the DRG 6 days after injury (48).

The homeostatic role of DRG macrophages, and the roles of macrophage infiltration and macrophage proliferation in the DRG in response to a peripheral injury have only begun to be explored (16, 47). Recently, CD163-macrophages were shown to be replenished more rapidly by circulating monocytes than CD163+ macrophages (47). Here, we found no differences in CD163+MHCII-macrophages after macrophage depletion in males or females with OA, and upregulated gene expression of *Cd163* in the DRG following depletion. Because this population of macrophages still expressed *Csf1r*, meaning that it was able to be targeted by the injection of AP20187, this suggests that this population replenished more rapidly than the other macrophage populations following depletion, potentially through local proliferation rather than through infiltration.

There were several limitations in this study. While we used two different time points in the male DMM model in addition to the female PMX model, we did not collect tissues at different time points following macrophage depletion. In addition, we assessed synovitis and F4/80+ cells in the joint, but we did not quantify changes in macrophage sub-populations in the joint. Finally, we only examined short-term effects on joint damage, but a strength of this study was that we intervened at time points when joint damage had already developed, which gave us some insight into the effects of targeting macrophages therapeutically.

### Conclusions

In conclusion, acute systemic macrophage depletion in mid-to late-stages of surgical models of OA alleviated pain-related behaviors and had no acute effect on joint damage severity in mice of both sexes. We gained mechanistic insights from DRG immune phenotyping, where M1-like MHCII+ macrophages were more affected by macrophage depletion than M2-like CD163+MHCII-macrophages in both sexes. RNA sequencing analyses revealed that the pro-inflammatory gene *Il1b* was consistently upregulated in the DRG with OA here and in other published datasets, and this gene was downregulated in the DRG with macrophage depletion. Future studies will continue to investigate the molecular mechanisms of DRG macrophages and how they contribute to persistent pain during OA progression.

## Declarations

### • Ethics approval and consent to participate

◦ All animal experiments were approved by the Rush University Institutional Animal Care and Use Committee.

### • Consent for publication

◦ Not applicable

### • Availability of data and materials

◦ The datasets used and/or analysed during the current study are available from the corresponding author on reasonable request.

### • Authors’ contributions

◦ Conceptualization: T.G., A.M.M., R.E.M. Investigation: T.G., S.I., A.M.O., J.L., L.W., N.S.A, H.L., F.K. Formal analysis: T.G., R.H., A.M.O., H.L., F.K., R.E.M. Visualization and data presentation: T.G., R.E.M. Writing: T.G. A.M.M., R.E.M. Reviewing and editing, T.G., A.M.M., R.E.M. Funding acquisition: T.G., A.M.M., R.E.M.

## • Acknowledgements

◦ We would like to thank the Rush University Flow cytometry core and the Comparative Research Center.

---

**Suppl. Fig. 1.**
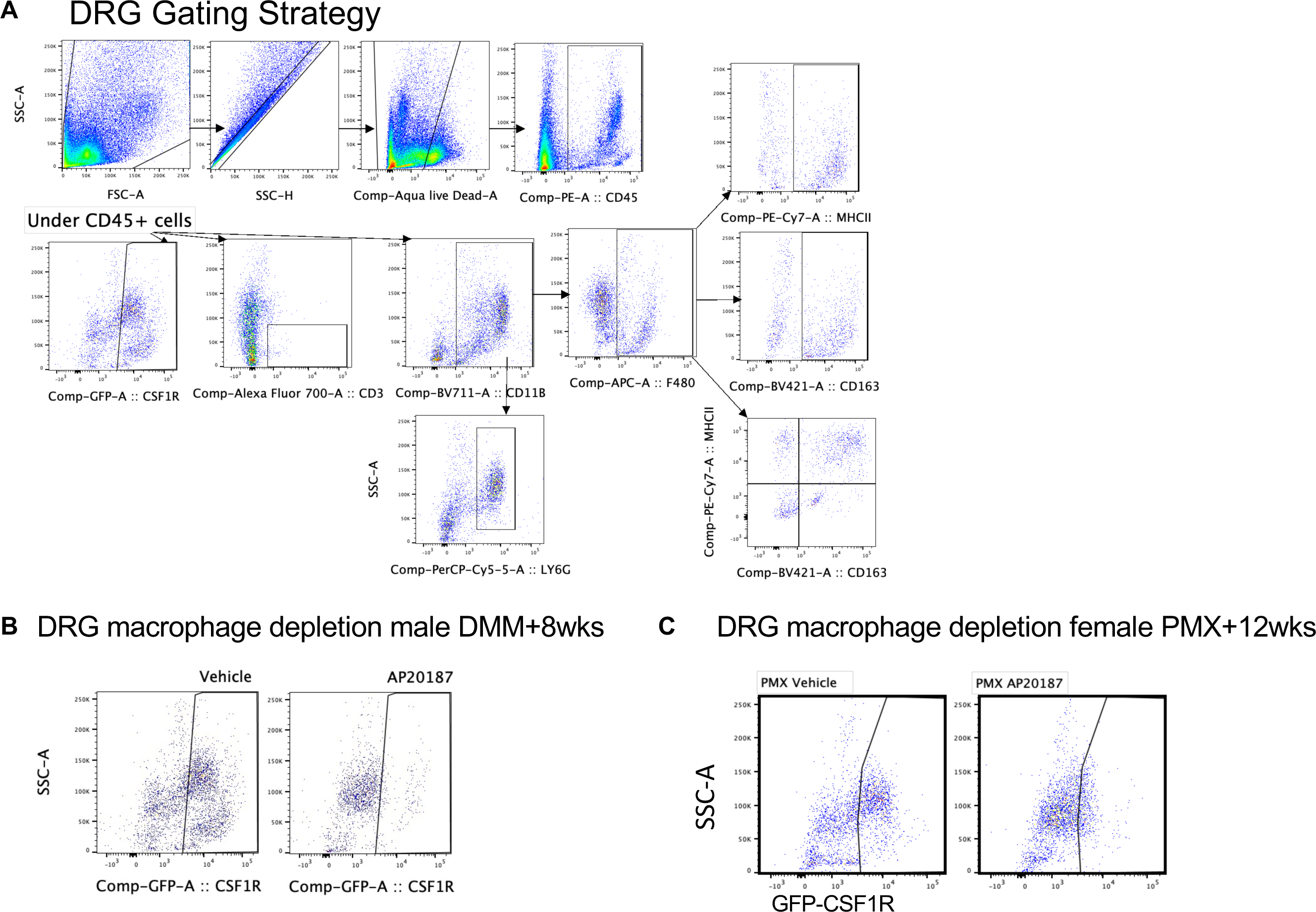
DRG Flow cytometry analysis. (A) DRG gating strategy. First all cells were gated for single cells using side-scatter height (SSC-H), then gated for live cells, then gated for CD45+ cells. Under CD45+ cells, CSF1R+ cells, CD3+ cells and CD11b+ cells were gated. Under CD11b+ cells, Ly6G+ neutrophils and F4/80+ macrophages were gated, and under F4/80+ cells MHCII and CD163 total positive cells were gated. Additionally, MHCII+ by CD163+ quadrant plots were made to evaluate the populations shown in Fig. 5. (B) Representative proof of CS1FR+ macrophage depletion in the DRG showing CSF1R+ cells in one Vehicle and AP20187-treated mouse from male MaFIA mice at 8 weeks after DMM surgery. (C) Same as in (B) for female MaFIA mice at 12 weeks after PMX surgery.

**Suppl. Fig. 2.**
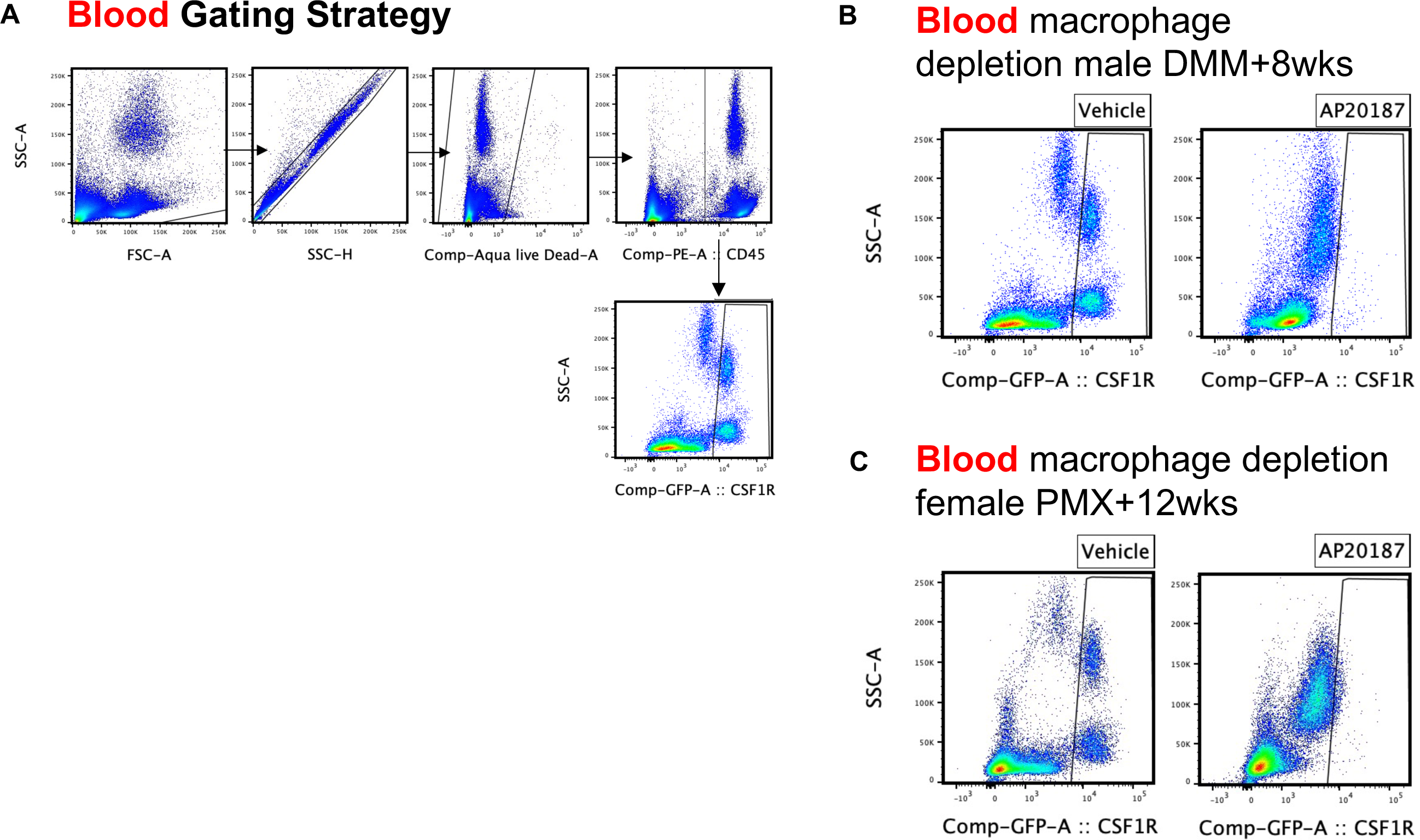
Peripheral blood flow cytometry analysis. (A) Gating strategy for peripheral blood. First all cells were gated for single cells using side-scatter height (SSC-H), then gated for live cells, then gated for CD45+ cells. Under CD45+ cells, CSF1R+ cells were gated. (B) Representative proof of CS1FR+ macrophage depletion in the blood showing CSF1R+ cells in one Vehicle and AP20187-treated mouse from male MaFIA mice at 8 weeks after DMM surgery. (C) Same as in (B) for female MaFIA mice at 12 weeks after PMX surgery.

**Suppl. Fig. 3.**
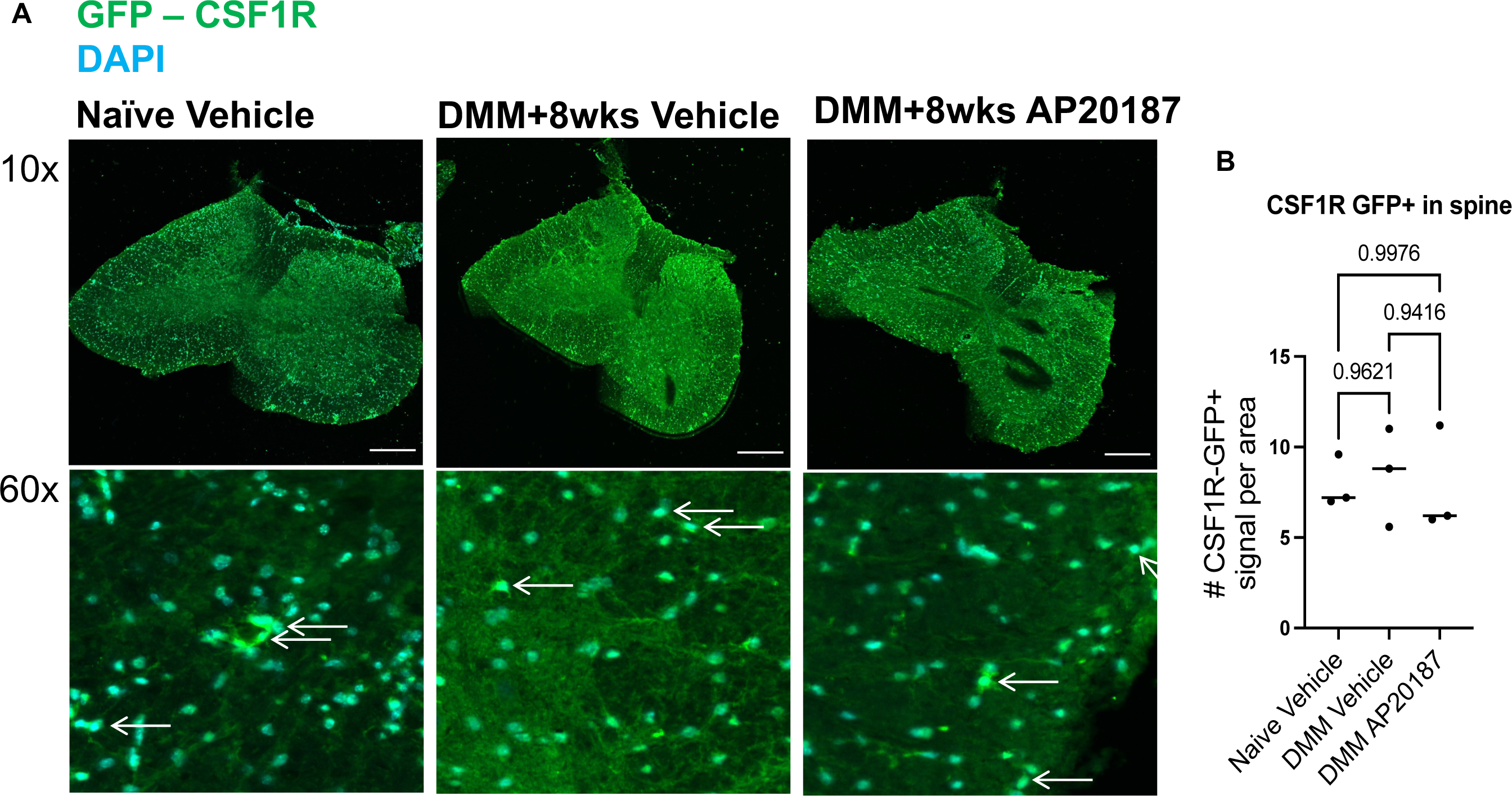
AP20187 does not deplete CSF1R+ cells in the dorsal horn. (A) Representative 10x and 60x magnification images of GFP+CSF1R+ cells in the dorsal horn of male MaFIA naïve mice given Vehicle, DMM mice at 8-weeks after surgery given Vehicle, or DMM mice at 8 weeks after surgery given AP20187. GFP in green, DAPI stained nuclei in teal/blue. White arrows point to GFP+CSF1R+ cells in 60x images. (B) Quantification of GFP+CSF1R+ cells in each experimental group. Scale bar is 30 μm. Statistical analysis by two-tailed t-test. Error bars show mean +/-95% CI. P values stated on graphs; significant if p < 0.05.

**Suppl. Fig. 4.**
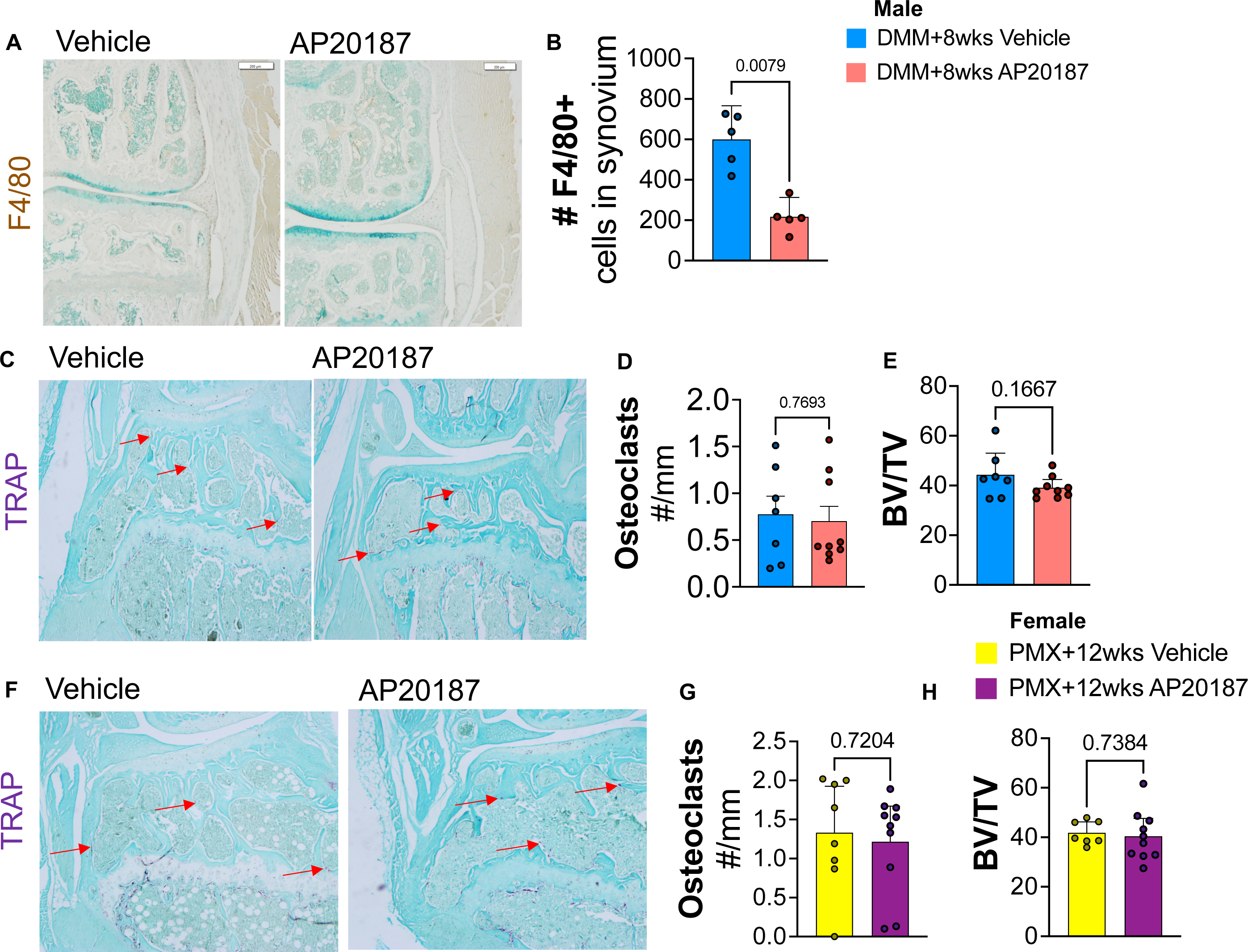
Decreased F4/80+ macrophages and no change in TRAP+ osteoclasts in knee joint with AP20187 macrophage depletion. (A) Representative 2x images of F4/80 stained knee joints from male MaFIA mice at 8-weeks after DMM surgery given Vehicle or AP20187 treatment. Brown signal represents F4/80+ signal. (B) Quantification of F4/80+ cells in knee synovium after treatment with Vehicle (n=5) or AP20187 (n=5) in male MaFIA mice at 8-weeks after DMM surgery. (C) Representative 4x images of TRAP+ stained knee joints from male MaFIA mice at 8-weeks after DMM surgery treated with Vehicle or AP20187. Red arrows point to positive TRAP signal in dark purple. (D) Quantification of number of TRAP+ osteoclasts per mm area and (E) Bone volume / Trabecular volume (BV/TV) ratio from male MaFIA mice knee joints at 8-weeks after DMM surgery treated with Vehicle (n=7) or AP20187 (n=9). (F) – (H) Same as in (C) – (E) but for female MaFIA mice knee joints at 12 weeks after PMX surgery given Vehicle (n=7) or AP20187 (n=9). Statistical analysis by two-tailed t-test. Error bars show mean +/-95% CI. P values stated on graphs; significant if p < 0.05.

**Suppl. Fig. 5.**
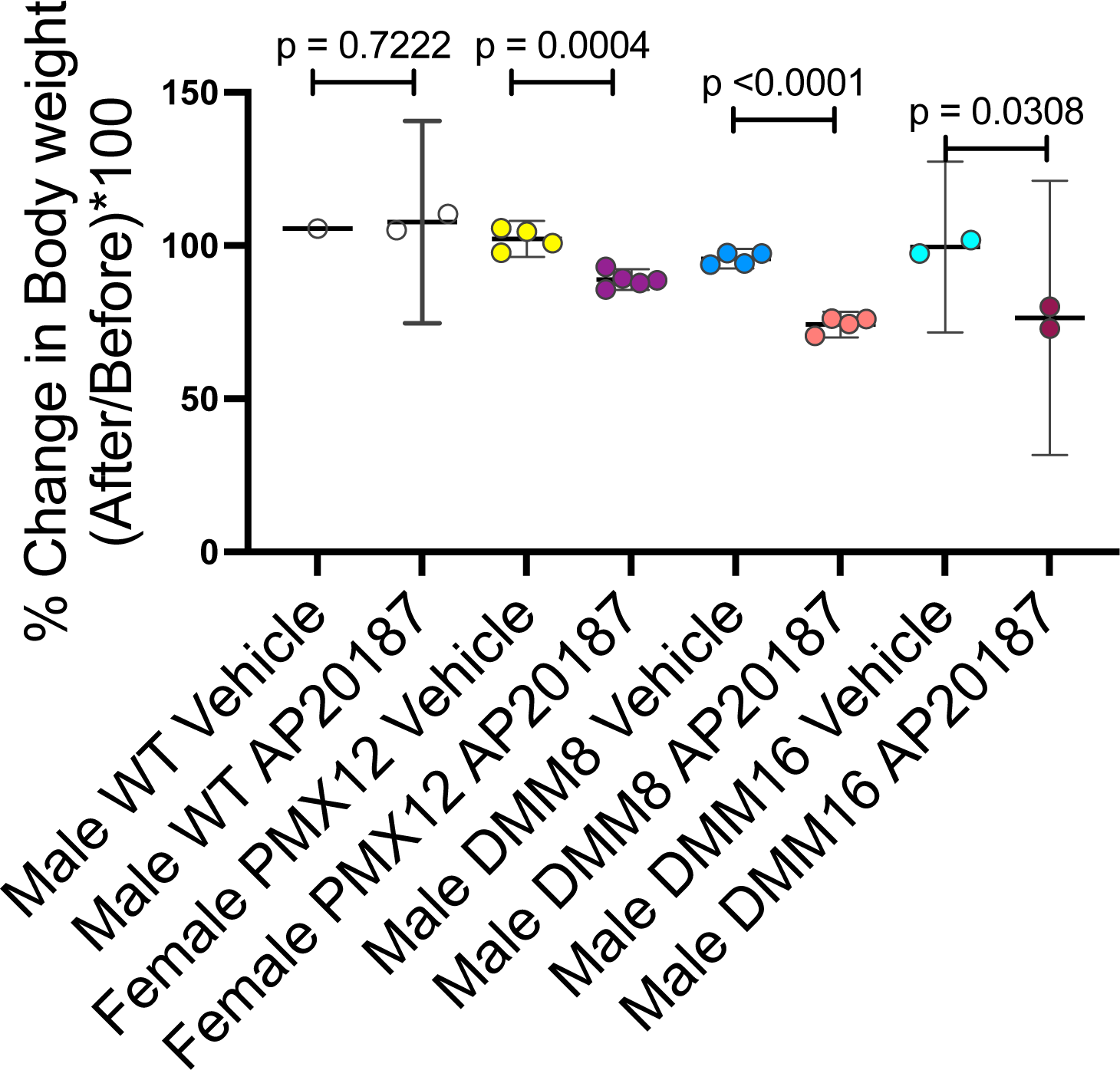
Decreased bodyweight in MaFIA mice treated with AP20187. Male wildtype (WT) mice treated with Vehicle (n=1) or AP20187 (n=2); female MaFIA mice at 12 weeks after PMX surgery were given Vehicle (n=4) or AP20187 (n=5); male MaFIA mice at 8 weeks after DMM surgery were given Vehicle (n=4) or AP20187 (n=4); and male MaFIA mice at 16 weeks after DMM surgery were given Vehicle (n=2) or AP20187 (n=2) for 5 consecutive days i.p. and mouse total bodyweight was measured before and after treatment. Statistical analysis by unpaired two-tailed t-test. Error bars show mean +/-95% CI. P values stated on graphs; significant if p < 0.05.

