## Supplementary material for "Acute systemic macrophage depletion in osteoarthritic mice alleviates pain-related behaviors and does not affect joint damage": Supp Table 1

**Supplementary Table 1.** List of antibodies used for flow cytometry analysis.

| **Antigen** | **Clone** | **Fluorophore** | **Source** |
| --- | --- | --- | --- |
| CD45 | 30-F11 | PE | BioLegend |
| CD3 | 145-2C11 | AF700, BV421 | BioLegend |
| CD11b | M1/70 | BV711 | BioLegend |
| MHCII (I-A/I-E) | M5/114.15.2 | PE/Cy7 | BioLegend |
| Ly6G | 1A8 | PerCP/Cy5.5 | BioLegend |
| Ly6C | HK1.4 | BV605 | BioLegend |
| F4/80 | BM8 | APC | BioLegend |
| CD8a | 53-6.7 | AF700 | BioLegend |
| CD4 | RM4-5 | PE/Cy7 | BioLegend |
| CD163 | S15049I | BV421 | BioLegend |
| CCR2 | SA203G11 | FITC, BV605 | BioLegend |
| Live/Dead | N/A | Aqua -405nm | ThermoFisherScientific |
